# Discovery of DTX3L inhibitors through a homogeneous FRET-based assay that monitors formation and removal of poly-ubiquitin chains

**DOI:** 10.1101/2023.03.13.532453

**Authors:** Carlos Vela-Rodríguez, Ilaria Scarpulla, Yashwanth Ashok, Lari Lehtiö

**Author notes:** Corresponding author: Lari Lehtiö –. Department of Genome Sciences, University of Washington, Seattle, USA.

## Abstract

Ubiquitination is a complex and reversible protein post-translational modification in which the subsequent action of enzymes belonging to three different families, broadly referred to as E1, E2 and E3, results in the covalent linking of ubiquitin to a target protein. While this linkage is canonically an isopeptide bond between the C-terminus of ubiquitin and the lysine residue of the target protein, Ser, Thr, and Tyr can also be susceptible to ubiquitination through an oxyester bond. Once ubiquitinated, multiple units of ubiquitin can be attached to the initial ubiquitin thus extending it to a chain of ubiquitins. Ubiquitination regulates multiple cellular processes, but it is best known as a modification that targets proteins for proteasomal degradation following the formation poly-ubiquitin chains linked through lysine 48 or 63 of ubiquitin. Dysregulation of ubiquitination has been associated with multiple types of cancer and efforts have been carried out to develop technologies that lead to the identification of inhibitors of the enzymes involved in the ubiquitination cascade. Herein, we present the development of a FRET-based assay that allows us to monitor auto-ubiquitination of DTX3L, a RING-type E3 ubiquitin ligase. Our method shows a robust signal window with a robust average Z’ factor of 0.76. From a validatory screening experiment we have identified the first molecules that inhibit DTX3L with potencies in the low micromolar range. Additionally, we have expanded the system to study deubiquitinases such as USP28 that lead to reduction of FRET due to hydrolysis of fluorescent poly-Ub chains.

## Introduction

The attachment of ubiquitin (Ub) to a target protein is a post-translational modification that regulates a wide range of events in eukaryotic organism including directing proteins for proteasomal or lysosomal degradation, regulating kinase activity, protein migration, and modifying protein-protein interactions (Komander, 2009; Schwartz and Ciechanover, 2009; Lipkowitz and Weissman, 2011; Ciechanover, 2015). Protein ubiquitination is a tightly regulated enzymatic cascade that canonically depends on the consecutive action of three enzyme families referred to as E1 ubiquitin activating enzymes, E2 ubiquitin conjugating enzymes, and E3 ubiquitin ligases (Koegl *et al*., 1999; Baranes-Bachar *et al*., 2018). E1 enzymes activate ubiquitin through adenylation and its subsequent transfer to an E2 enzyme (Schulman and Harper, 2009; Hann *et al*., 2019). Primed E2s are recruited by Ub-E3 ligases, which simultaneously recognise the target protein and facilitate the discharge of Ub from the E2 to the substrate protein (Pruneda *et al*., 2012; Soss, Klevit and Chazin, 2013). E3 ligases are the most abundant members of the ubiquitination cascade with reports of even up to ∼700 putative ligases encoded in the human genome in comparison with 2 E1 enzymes and 40 E2 enzymes (Stewart *et al*., 2016). Depending on their transfer mechanism, E3s are further classified as RING, HECT, and RING-between-RING (RBR) (Yang *et al*., 2021).

DTX3L is a RING-type ubiquitin ligase characterised by a conserved C-terminal domain that mediates ubiquitination of target proteins and ADP-ribosylation of ubiquitin (Obiero, Walker and Dhe-Paganon, 2012; Yang *et al*., 2017; Ahmed *et al*., 2020; Chatrin *et al*., 2020). However, recent studies on DTX2, member of the same protein family as DTX3L, suggest that the product of DTX proteins is a hybrid chain resulting from the ubiquitination of ADP-ribose on already modified proteins (Zhu *et al*., 2022). *In vitro* studies identified that DTX3L formed K6-, K11-, K48- and K63-mediated poly-Ub chains (Ashok *et al*., 2022); supporting the role of DTX3L in proteasomal degradation and DNA damage response pathways (Yan *et al*., 2013).

DTX3L has recently gained attention as a potential drug target due to its involvement in anti-viral response and overexpression in various types of cancer (Vela-Rodríguez and Lehtiö, 2022). In the context of cancer it has been associated with lymphoma, multiple myeloma (MM), glioma, and prostate and breast cancer (Juszczynski *et al*., 2006; Bachmann *et al*., 2014; Thang *et al*., 2015; Shen *et al*., 2017; Xu *et al*., 2017; Yang *et al*., 2017; Tang *et al*., 2018). The specific roles that DTX3L plays in all these events are still rather obscure, but it has been shown that it regulates NOTCH signalling (Matsuno *et al*., 1995; Yamamoto *et al*., 2001; Zhang *et al*., 2010) and mono-ubiquitination of histone H4 by DTX3L leads to DNA protection resulting in the development of chemo-resistant cancer (Bachmann *et al*., 2014).

Ubiquitination can be removed by deubiquitinases (DUBs), ∼100 in the human genome, which are responsible for the hydrolysis of the isopeptide bond between Ub moieties (Clague, Urbé and Komander, 2019), thus recycling the pool of free Ub. DUBs are classified in the groups of cysteine proteinases and metalloproteases with distinct reaction mechanisms (Snyder and Silva, 2021). We have previously identified that at least Ubiquitin Specific Protease 28 (USP28) can remove the Ub chains generated by DTX3L both from DTX3L itself and from USP28 *in vitro* as well as in cellular context thereby regulating the DNA repair pathways (Ashok *et al*., 2023).

As ubiquitination is involved in several pathologies, there have been constant efforts in targeting the different components of the ubiquitination machinery and several assay technologies have been developed. These assays have made use of DELFIA, ELISA and FRET technologies but they are mostly limited to track auto-modification of the E3s or E2s due to the inherent complexity of the reaction. Many of the available assay technologies that study the ubiquitination cascade rely on protein immobilisation followed by multiple washing steps (Kenten *et al*., 2005; Ceccarelli *et al*., 2011; Rossi *et al*., 2014), or they require chemical labelling of Ub (Boisclair *et al*., 2000; Gururaja *et al*., 2005; Madiraju *et al*., 2012; Krist *et al*., 2016; Wu *et al*., 2016, 2021).

Here we describe the development of a homogeneous FRET-based assay that monitors in real time the formation and hydrolysis of poly-Ub chains demonstrated with the activities of DTX3L and USP28. The method uses fluorescent fusion proteins (CFP-Ub and YFP-Ub) to avoid labelling steps making the assay easy to adopt in different laboratories with minimal running costs. The system has been validated in 384-well plate format with a robust screening window coefficient Z’ value of 0.76 for both ubiquitination and deubiquitination reactions. The assay was further validated using a known USP28 inhibitor for potency measurement and in screening of a random compound library allowing us to describe the first small molecule inhibitors of DTX3L.

## Results

### Fluorescent ubiquitin exhibits native-like behaviour in the ubiquitination reaction

In order to track the formation of poly-Ub chains we produced fusion proteins of Ub with mCerulean and mCitrine to generate donor and acceptor fluorophores, hereinafter referred to as CFP-Ub and YFP-ub, respectively (**Figure 1A**). Initial expression trials of these fluorescent constructs produced inclusion bodies, which was overcome by increasing the length of the flexible linker between the fluorescent protein and Ub to 16 residues, including a tobacco etch virus (TEV) recognition site (**Supplementary Figure S1)**. We also confirmed through SEC-MALS that the fusion proteins behave as monomers in solution, since the fluorescent proteins used in the study are enhanced, monomeric variants of CFP and YFP (**Supplementary Figure S2**). Using a gel-based ubiquitination assay reported by us earlier (Ashok *et al*., 2022), we verified that the incorporation of the fluorescent proteins to the N-terminus of ubiquitin did not interfere with the auto-ubiquitination reaction of DTX3L (**Figure 1B, lanes 6 and 8**). Electrophoresis of the samples that have not been subjected to heat denaturation also showed that the formation of poly-ubiquitin chains does not affect the fluorescence of CFP nor YFP (**Figure 1B, lower panel lanes 12** & **14**). The assay was set up using a fragment of DTX3L that lacks the N-terminal domains and can be easily produced in *E. coli* (Ashok *et al*., 2022). These initial trials with a gel-based assay indicate that fluorescent fusion ubiquitin is compatible with the ubiquitination reaction in a native-like manner, allowing the development of a FRET-based assay by exploiting DTX3L auto-modification mechanism with the proposed schematic showed in **Figure 1C**.

**Figure 1.**
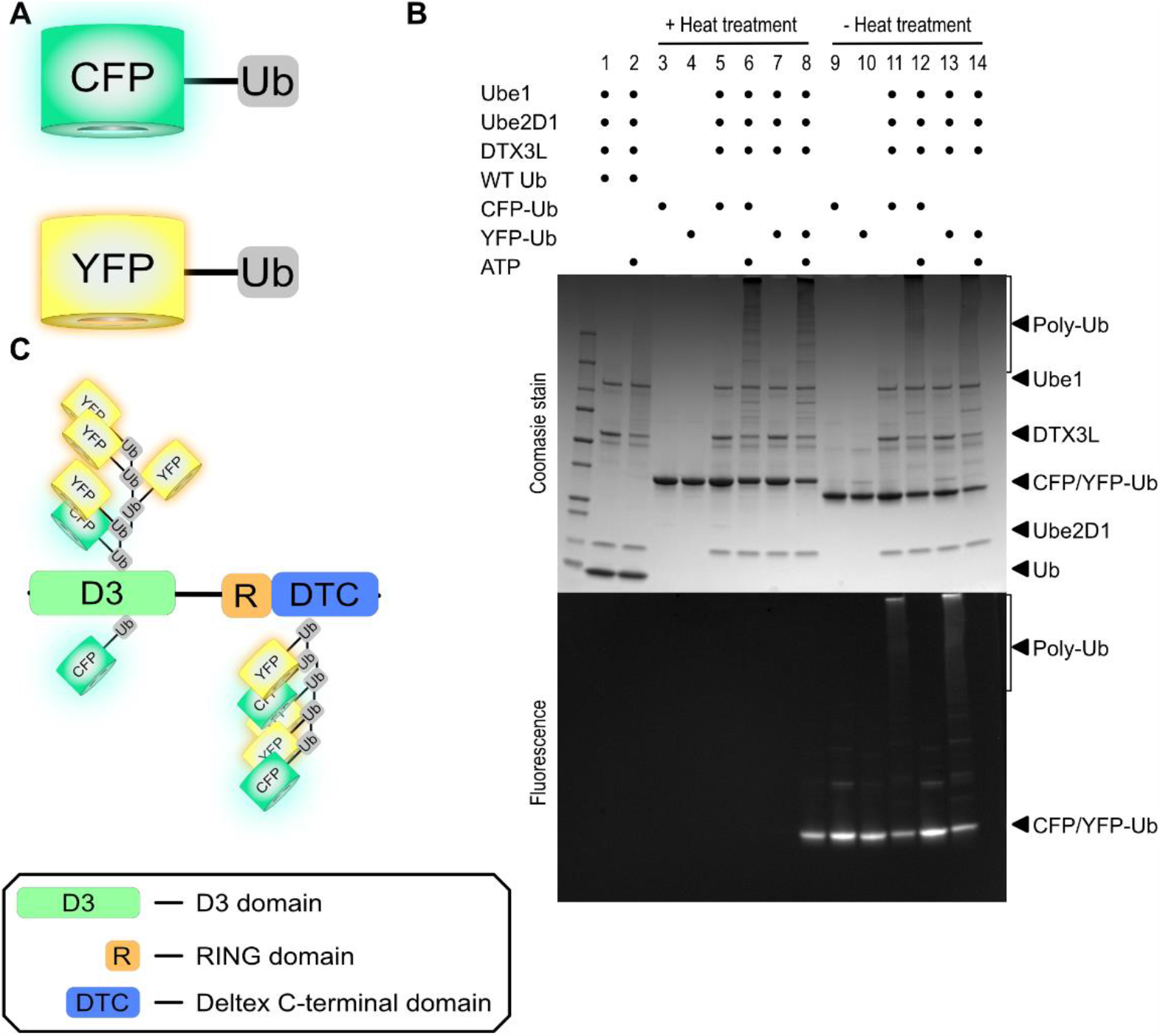
Fluorescent ubiquitin fusion proteins can be used in a ubiquitination assay. A) Schematic of the fluorescent ubiquitin constructs used in the assay. B) SDS-PAGE analysis of ubiquitination reaction confirms that DTX3L accepts the fluorescent fusion proteins as substrate for the formation of poly-Ub chains in a native-like manner (upper panel). The formation of poly-Ub chains does not affect the fluorescence of the chromophores (lower panel). C) Schematic of the FRET-based ubiquitination assay. In a reaction including Ube1 (E1), Ube2D1 (E2), DTX3L (E3) and ATP, DTX3L undergoes self-modification, and the formation of poly-ubiquitin chains translates in energy transfer from CFP-Ub to YFP-Ub.

### Fluorescent ubiquitination produces a robust signal that allows identification of DTX3L inhibitors

During the development of the assay, we opted for a ratiometric approach in which the signal was obtained by dividing the fluorescence recorded at 477 nm (emission peak for mCerulean) by the fluorescence recorded at the 527 nm (emission peak for mCitrine). Ratiometric FRET (rFRET) was used over FRET efficiency, or quenching due to its relative simplicity, and the little effect that small variations derived from dispensing or multiwall plates have on the measurement. In the gel-based assay (**Figure 1B**), we have used the absence of ATP to prevent ubiquitination from proceeding as ubiquitin cannot be activated by Ube1. However, we intended that the assay would focus on the activity of DTX3L, and the omission of ATP in the reaction affects upstream enzymes in the ubiquitination chain. We therefore generated a DTX3L double mutant (DTX3L^CS^) where C561 and C564 (C1 and C2 of the Zinc-coordinating RING domain) were mutated to S as the mutant has been previously reported to lack Ub E3 ligase activity (Zhang *et al*., 2015). We confirmed that the mutation had a minimum effect on the folded state of the protein (**Supplementary Figure S3**) and that the construct lacked E3 ligase activity. This loss of activity was observed as the absence of the characteristic poly-Ub smear in SDS-PAGE (**Supplementary Figure S4**) as well as the low rFRET signal in our initial trials with DTX3L (**Figure 2B and Supplementary Figure S5**). The behaviour of the mutant would remain consistent throughout the development of the assay, indicating that the low signal recorded with DTX3L^CS^ was not derived from poor reaction conditions (**Figure 2B**). As our starting point, we scaled down the protein concentration for Ube1 and Ube2D1 from the original gel-based assay to 12 nM and 60 nM, respectively. As E1s are the enzymes with the quickest turnover in the ubiquitination cascade (Jin *et al*., 2007), we opted to have it as the enzyme with the lowest concentration. On the other hand, as our system makes use of the auto-modification of DTX3L, thus acting both as the ligase and the substrate, we set the concentration to 500 nM. Additionally, we first screened the reaction at different equimolar concentrations of CFP-Ub and YFP-Ub, where the rFRET was measured at different incubation times (**Figure 2A**). Upon noticing that the rFRET at 2 h incubation and a fluorophore concentration of 40 nM (**Figure 2A, open circles**) gave a reasonable signal, we undertook a series of steps to improve the assay conditions. During the optimisation stage, we varied the fluorophore ratio, the ratio between Ube1 and Ube2D1, buffer composition, and the final concentration of ATP in the reaction (**Figure 2B and Supplementary Figure S6**). At the end of an iterative optimisation process, we were able to identify conditions in which the rFRET of the reaction with DTX3L was three times higher than with DTX3L^CS^ (**Figure 2B**). Furthermore, the signal was saturated already at 80 min of incubation and sufficient signal was achieved already at 40 min (**Supplementary Figure S7**). The final assay conditions were defined as 12 nM Ube1, 180 nM Ube2D1, 500 nM DTX3L, 25 nM CFP-Ub and 350 nM YFP-Ub. Enzymes were mixed in 10 mM HEPES (pH 7.5) and the reaction was initiated by the addition of ubiquitination buffer [Tris (5 mM, pH 7.5), ATP (0.2 mM), MgCl_2_ (0.5 mM) and DTT (0.2 mM)].

**Figure 2.**
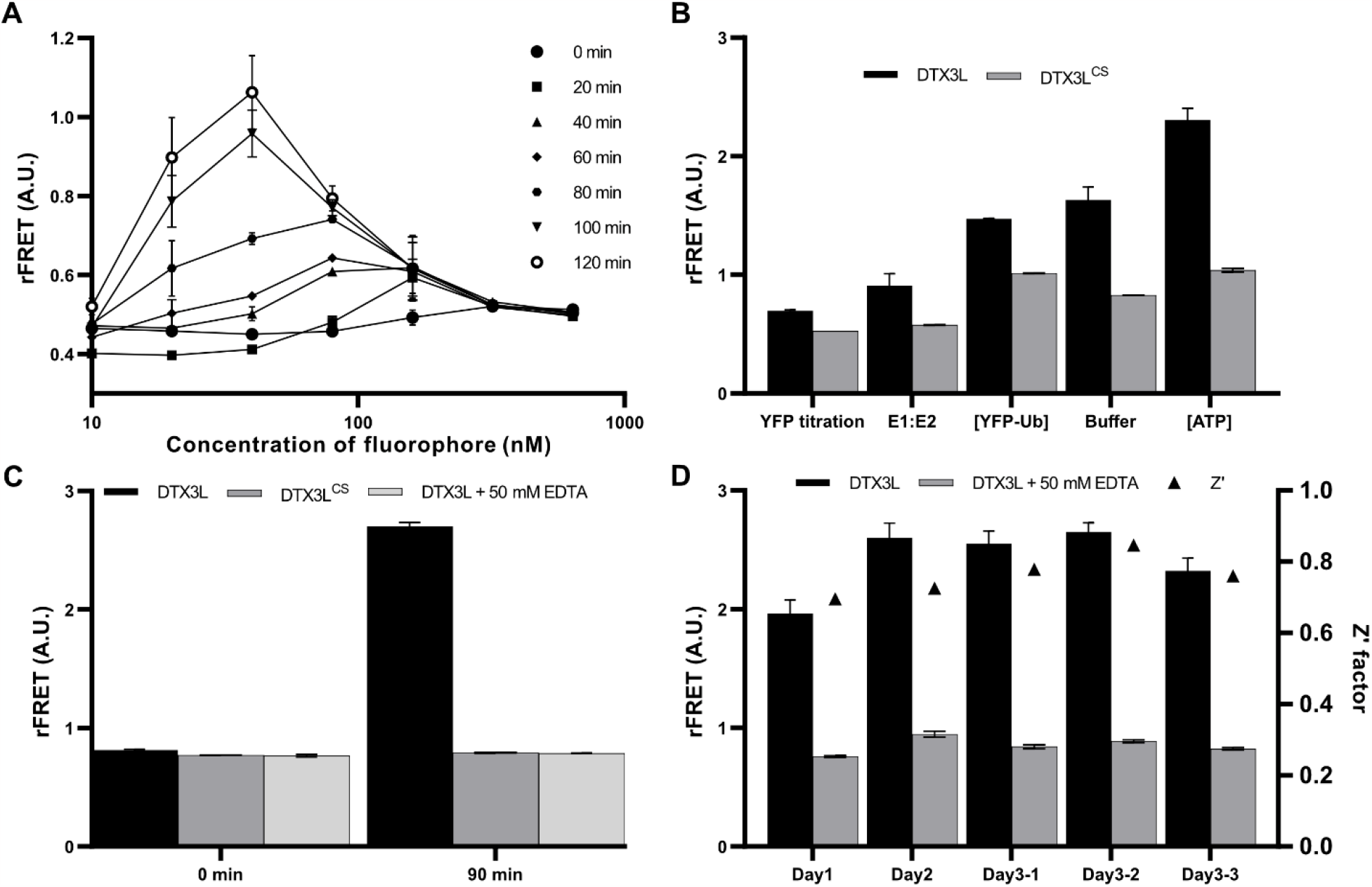
Optimisation of the assay conditions generates a robust signal that is suitable for high-throughput screening based on its statistical validation. A) Effect of fluorophore concentration on the rFRET of the ubiquitination assay. For the experiment, CFP-Ub and YFP-Ub were mixed at equimolar concentrations. B) Summary of the individual steps undertaken in the optimisation of the assay. The results are displayed in the progressive manner that they were performed, comparing the signal of DTX3L to that of DTX3L^CS^. C) Addition of 50 mM EDTA to a reaction with DTX3L reproduces the signal obtained using DTX3L^CS^. The signal was measured immediately after the addition of ATP (0 min) and after an incubation of 90 min. D) Statistical validation of the assay indicates suitability for the discovery of small molecule inhibitors. Data shown in panels A-C correspond to mean ± SD of four replicates. Data shown in panel D corresponds to mean ± SD of 176 replicates per plate.

For the inhibitor screening we aimed to have a condition that mimics chemical inhibition of DTX3L. Since the lack of activity of DTX3L^CS^ is derived from the disruption or the RING finger, we deduced that using a chelating agent to sequester Zn^2+^ from DTX3L would abolish ligase activity. We adopted the use of EDTA as it has been previously showed that it can stop the ubiquitination of p53 by ICP0, another RING-type Ub-E3 ligase (Canning *et al*., 2004). Reactions in which EDTA was added immediately after the addition of ATP showed similar signal as DTX3L^CS^ (**Figure 2C**). Subsequently, in the lack of DTX3L inhibitors, we used reactions with EDTA as positive controls and reactions without EDTA as negative controls for the evaluation of the assay performance. The validation of the assay gave a Z’ factor with a value in a range from 0.69 to 0.84 (average Z’ factor = 0.76) with a significant signal window (**Figure 2D and Supplementary Table S1**). Statistical validation supported the suitability of the assay for the screening of small molecules that affect the progression of the ubiquitination reaction.

### FRET-based ubiquitination assay leads to the discovery of DTX3L inhibitors

To test the assay performance, we screened three libraries from the National Cancer Institute (NCI) to identify small molecules that interfere with the ubiquitination reaction (**Figure 3**). As the compounds are dissolved in DMSO, we first performed a DMSO tolerance test to evaluate the effect of DMSO on DTX3L auto-ubiquitination. The DMSO tolerance test indicated that our system performs well with concentrations of DMSO to up to 7.5% (**Figure 3A**). As DTX3L is a RING-type E3 ligase there are no discernible pockets that might bind small molecules and there are also no known inhibitors reported for it. We therefore carried out the screening at a high concentration of 100 μM (1% DMSO) to identify initial hit compounds from a total of 2428 compounds (**Figure 3B**).

**Figure 3.**
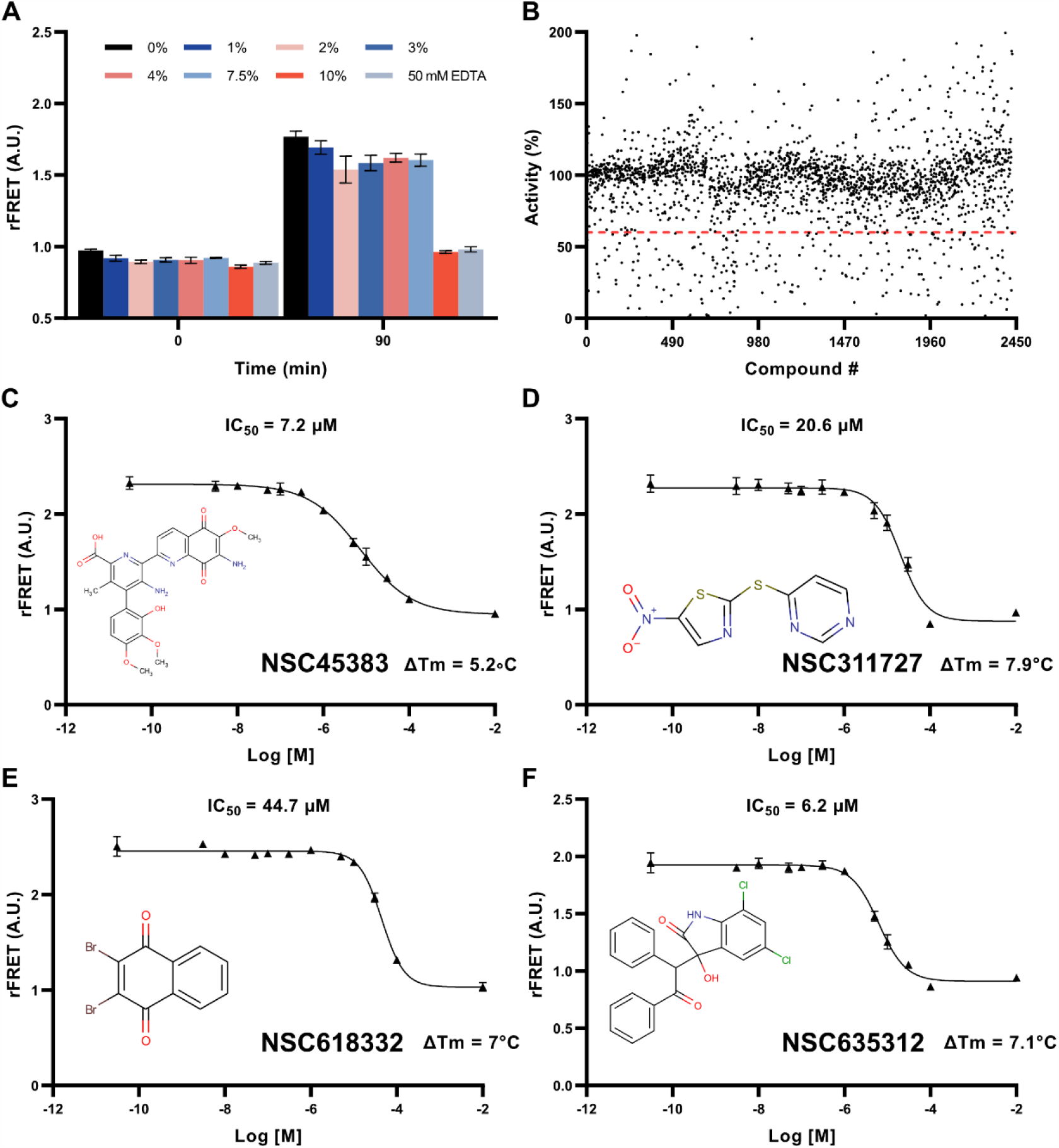
Screening of three small molecule libraries identifies compounds that inhibit the progression of the ubiquitination reaction by binding to DTX3L. A) DMSO tolerance test indicates that the FRET ubiquitination is not affected by the addition of DMSO to even up to 7.5%. B) Initial screening by single measurements detects several compounds that show inhibition towards the ubiquitination reaction. Dots below red dotted line shows hits C-F) IC_50_ curves for the selected compounds with the chemical structure of the compound and the differential melting temperature (ΔTm) calculated using nanoDSF. IC_50_ values indicate modest potencies, and positive shifts in the ΔTm of DTX3L confirm binding of the compounds to the protein. Data shown in panel A correspond to mean ± SD of four replicates. IC_50_ curves shown in panels C-F corresponds to mean ± SD of 4 replicates. ΔTm values corresponds to the mean of 3 replicates.

During the screening process we used 410 and 430 nm as excitation wavelengths under the reasoning that some compounds may exhibit fluorescence at one wavelength or the other, strategy that has been used successfully in previous studies (Sowa *et al*., 2020). From the primary screening, we selected as potential hits only those molecules that showed an apparent inhibition higher than 40% with less than 10% difference in inhibition between excitation wavelengths. Furthermore, the compounds selected had to show an increase in the emission at 477 nm and a decrease in the emission at 527 nm as compared to the average values of the negative control. Our reasoning for this constraint was that for the compound to inhibit DTX3L, therefore the formation of poly-Ub chains, the fluorescence of YFP (527 nm) is reduced as an effect of a reduced excitation that translates in fluorescence of CFP (477 nm) being re-established.

Based on these filters, we selected 50 compounds that were re-evaluated at 10 μM, from which we selected five compounds that showed an apparent inhibition higher than 20% using 410 and 430 nm excitation wavelength. For the selected compounds, we measured the IC_50_ values with 100 nM starting concentration to confirm the inhibition of the ubiquitination reaction (**Figure 3C-F**). Based on this evaluation we ruled out NSC265211 as a false positive as inhibition was not detected. We used the gel-based ubiquitination assay as an orthogonal method to assess the effect of the selected compounds on DTX3L auto-ubiquitination, judged by the intensity of the smear (**Supplementary Figure S8**). Subsequently, as we were interested in finding compounds that affect DTX3L, we evaluated the direct binding of the four remaining compounds to DTX3L through differential scanning fluorimetry (DSF) (**Supplementary Table S2**). Through this evaluation, all compounds showed positive shifts in the melting temperature hinting that they directly bind to DTX3L. However, NSC311727 and NSC618332 showed a decrease in the onset temperature for scattering as compared to the control sample, indicating that they might promote protein aggregation.

Compound NSC45383 is commonly known as streptonigrin (**Figure 3C**). Streptonigrin is an antibiotic isolated from *Streptomyces flocculus* that induces DNA breaks and while previously been used in cancer therapy, its cytotoxic effects led to a decrease in its usage (Bolzán and Bianchi, 2001). In a more recent study it was showed that the toxicity is dependent on the concentration and that streptonigrin prevents STAT3 phosphorylation (Loyola *et al*., 2020). The activity of NSC311727 has been assessed by the Development Therapeutics Program (DTP) of the NCI and observed to inhibit cell growth in at least 90% on multiple assays that screened growth inhibition of different cell lines of non-small cell lung cancer. Similarly, NSC618332 and NSC635312 were classified as potentially bioactive against cancer as they inhibited growth of yeast strains that bore alterations in the DNA damage repair or cell cycle regulation.

### Detection of the poly-Ub chains hydrolysis

We have previously reported that there is a mutual regulation between USP28 and DTX3L in which the DUB hydrolyses poly-Ub chains generated by DTX3L. Ubiquitination of USP28 by DTX3L was also demonstrated with a catalytically inactive USP28^C171A^ mutant (Ashok *et al*., 2023). We therefore sought to test if addition of USP28 to pre-formed poly-Ub chains on DTX3L would lead to reduction in rFRET due to hydrolysis poly-Ub chains. As addition of EDTA leads to inactivation of DTX3L, we hypothesised that addition of EDTA after a fixed time period would arrests further ubiquitination.

To test our hypothesis, we prepared ubiquitination reactions with CFP-Ub and YFP-Ub and after incubating for 150 min we added 50 mM EDTA. Simultaneously, USP28 and a catalytically inactive mutant USP28^C171A^ were added to the reactions (**Figure 4A**). We noticed that addition of EDTA decreased the rFRET through an unknown mechanism. Despite this decrease, reactions lacking USP28 (Ub. arrested) and reactions with USP28^C171A^ showed very similar values. On the other hand, arrested reactions with USP28 showed a progressive decrease in the rFRET, confirming that USP28-mediated hydrolysis can be monitored with our system.

**Figure 4.**
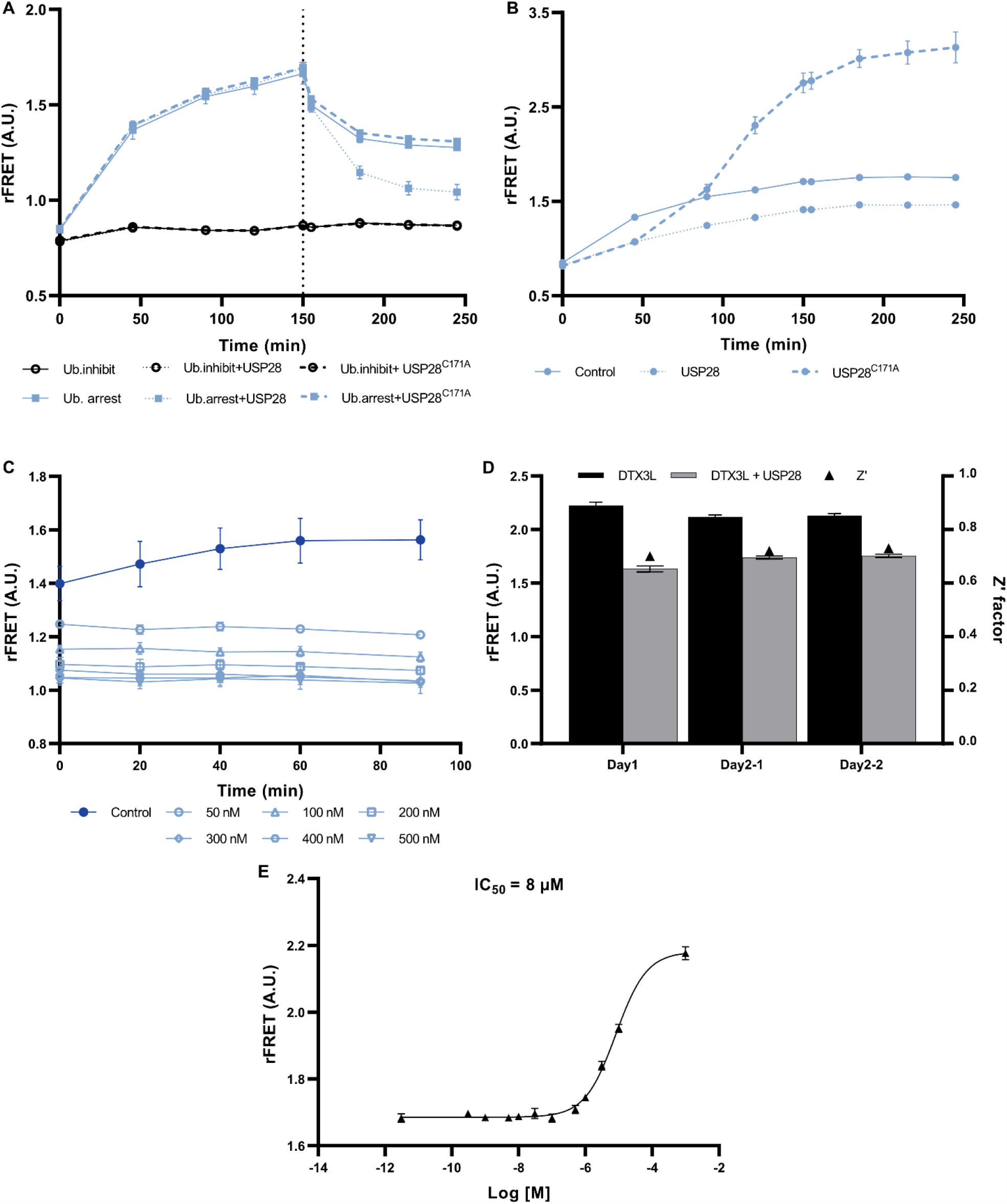
Digestion of poly-Ub chains by USP28. A) Addition of USP28 and USP28^C171A^ to inhibited ubiquitination reactions and ubiquitination reactions arrested after 150 min with 50 mM EDTA. B) Addition of USP28 and USP28^C171A^ to ubiquitination reactions. C) Titration of the ubiquitination reaction with different USP28 concentrations. D) Plate validation of the FRET assay to monitor hydrolysis of poly-Ub chains. E) IC_50_ measurement of AZ1 with the FRET ubiquitination assay. In addition to USP28 or USP28^C171A^, all reactions contain Ube1, Ube2D1 and DTX3L. Data shown in panels A-C and E correspond to mean ± SD from four replicates whereas in panel D there are 176 data points per plate.

We then assessed the effect of incorporating USP28 and USP28^C171A^ to the ubiquitination reactions from the beginning and in absence of EDTA (**Figure 4B**). Ubiquitination reactions to which USP28 was added showed an rFRET about 50% lower than the control reactions, likely due to an equilibrium between the writing and erasing steps. Up to 45 min USP28^C171A^ seems to have the same effect as USP28. However, at 90 min reactions with USP28^C171A^ have an rFRET comparable to that of the control group and at any time point after that, the rFRET is higher than the control. Since USP28^C171A^ acts as a substrate for DTX3L, the activity of the ligase is not limited to auto-modification, resulting in a boosted activity.

We also evaluated the effect of USP28 and USP28^C171A^ in inhibited ubiquitination reactions (**Figure 4A**), noticing that the deubiquitinase does not have an inherent effect on the rFRET.

Having verified that FRET can be used to monitor the poly-Ub chains hydrolysis by USP28, we measured the rFRET using different concentrations of USP28 to determine the conditions used in a de-ubiquitination assay (**Figure 4C**). Our observations suggested that the rFRET decreases with increasing concentrations of USP28. This can be translated in a more effective hydrolysis, which is not perceived after 300 nM of USP28 (**Figure 4C, open rhombus**). We observed that by using 50 nM USP28 (**Figure 4C, open circle**) we were able to decrease the rFRET considerably with very little variation and through the statistical validation using that concentration (**Figure 4D and Supplementary Table S3**) we determined that the signal window was enough to determine an average Z’ value of 0.7.

As a final confirmation that the assay was able to detect inhibition of USP28, we measured the IC_50_ value of a known dual inhibitor for USP28 and USP25 (AZ1) using the FRET-based assay (**Figure 4E**). We previously showed that in cells this compound is able to affect DTX3L in a similar fashion as knock down of USP28 (Ashok *et al*., 2023) and the IC_50_ value determined with our method (IC_50_ = 8 μM) is close to the binding affinity (K_d_ value of 3.7 μM) measured previously with microscale thermophoresis (MST) (Wrigley *et al*., 2017). This indicates that our system can be adopted to identify and characterize inhibitors also for the DUB.

## Discussion

Ubiquitin signalling is a potential drug target in cancer therapies and therefore efforts have been made to develop assays amenable for high-throughput screening of inhibitors affecting the enzymes involved in this cascade (Morrow *et al*., 2015; Fricker, 2020; Aliabadi *et al*., 2021). Perhaps the most limiting factor in assay development for the ubiquitin system is the lack of a universal consensus sequence as opposed to Ub-like modifications (Sampson, Wang and Matunis, 2001). Despite some ubiquitination motifs having been identified and described in the literature (Jadhav and Wooten, 2009; Davey and Morgan, 2016), most assay technologies rely on the auto-modification of either the E2 or the E3 enzymes (Gururaja *et al*., 2005; Kenten *et al*., 2005; Ceccarelli *et al*., 2011; Madiraju *et al*., 2012; Rossi *et al*., 2014; Krist *et al*., 2016). Only a few described assays do not rely on the auto-modification, but rather make use of natural substrates (Xu *et al*., 2005; Wu *et al*., 2021). Herein, we have described an assay capable of identifying potential inhibitors of DTX3L. The assay relies on the auto-modification of DTX3L making it easier to carry out (**Figure 2**) but we also show that it is also compatible with a natural a substrate that helps to enhance the signal, resulting in an increased sensitivity if needed (**Figure 4A**).

While fluorescent proteins have been used in the assay development for Ub-like enzymatic cascades (Bossis *et al*., 2005; Stankovic-Valentin *et al*., 2009; Song and Liao, 2012; Malik-Chaudhry, Saavedra and Liao, 2014; Wiryawan *et al*., 2015), the approach has not yet been explored in ubiquitination. The use of fluorescent fusion tags bypasses labelling steps and avoids the generation of site-specific mutations that facilitate the chemical labelling of ubiquitin, thus streamlining the process. Additionally, omission of labelling or immobilisation steps can considerably reduce assay times. One of the limitations of the assay in its current state is that the expression of a fusion CFP/YFP-Ub prevents the formation of linear poly-Ub chains through the N-terminal amide. This type of chain is only mediated by the linear ubiquitin chain assembly complex (LUBAC) (Kirisako *et al*., 2006; Gerlach *et al*., 2011; Ikeda *et al*., 2011; Tokunaga *et al*., 2011; Fujita *et al*., 2018), which is why expressing fusion proteins should be compatible with the majority of the Ub E3 ligases and E2 conjugating enzymes. Most of the available assay technologies designed to study poly-ubiquitination reactions block the N-terminus achieving comparable results as the ones we have obtained (Boisclair *et al*., 2000; Madiraju *et al*., 2012; Wu *et al*., 2021).

DTX3L seems to have multiple functions and to date its role in cancer biology is not completely clear (Vela-Rodríguez and Lehtiö, 2022). This correlates to reported evidence for other E3 ligases, e.g. FBXW7 and βTrCP, that act as tumour suppressors or as oncogenic proteins depending on the context (Saitoh and Katoh, 2001; Mao *et al*., 2004; Ougolkov *et al*., 2004; Kim *et al*., 2007; Matsuoka *et al*., 2008; Busino *et al*., 2012). In a similar way, USP28 has been reported to act both as a positive and negative regulator in cancerous cells even though it was initially considered to be a strict oncoprotein (Prieto-Garcia *et al*., 2021). Both DTX3L and USP28 seem to regulate ligation and hydrolysis, respectively, of K6-K11-, K48- and K63-linked Ub-chains (Zhen *et al*., 2014; Ashok *et al*., 2022). Poly-ub chains linked through K48 mainly adopt a closed conformation forming a hydrophobic core between two Ub moieties; K63-mediated chains on the other hand prefer open conformations (Li and Ye, 2008). This linkage preference of DTX3L and USP28 supports the viability of our system with chains of both conformations.

While there are inhibitors of USP28 like the one used in this study (Wrigley *et al*., 2017), there are no known inhibitors for DTX3L or any other members of its family. In our work, we describe the initial inhibitor for the Ub-ligase function of DTX3L that despite the low potency, have a considerable effect on the formation of poly-Ub chains. The assay that we have developed can potentially be adapted to test the activity of Ub E3 ligases that undergo auto-modification with a suitable E2 enzyme or if there is a target protein available and identify not only potential inhibitors of the ubiquitination cascade but also possible enhancers of the modification.

## Materials and methods

### Cloning

Unless stated otherwise, coding sequences of the proteins of study were cloned into expression vectors by sequence and ligation independent cloning (SLIC) (Jeong *et al*., 2012), using *E. coli* NEB5α for cloning procedures. The expression vectors pNIC28-MBP, pNIC-CFP and pNIC-YFP were generated by the insertion of MBP (Sowa *et al*., 2020), mCerulean and mCitrine between the His6-tag and TEV protease recognition site of pNIC28-Bsa4 (Sowa *et al*., 2020, 2021). Codon optimised sequence for USP28 was procured from Genescript and cloned between the NdeI/BamHI multiple cloning site (MCS) of pMJS162 (Gaciarz *et al*., 2016). Fluorescent ubiquitin was generated by the insertion of the coding sequence of ubiquitin into pNIC-CFP or pNIC-YFP. Generation of the inactive DTX3L mutant (DTX3L^CS^) was achieved by site-directed mutagenesis in a protocol adapted to polymerase chain reaction (PCR) (Edelheit, Hanukoglu and Hanukoglu, 2009). All constructs were verified by dideoxy sequencing. Details about the expression constructs are shown in **Supplementary Table S4**. Ube1/PET21d was a gift from Cynthia Wolberger (Addgene plasmid # 34965; http://n2t.net/addgene:34965; RRID:Addgene_34965) (Berndsen and Wolberger, 2011). UbcH5A was a gift from Cynthia Wolberger (Addgene plasmid # 61081; http://n2t.net/addgene:61081; RRID:Addgene_61081) (Wiener *et al*., 2013). Ubiquitin WT was a gift from Rachel Klevit (Addgene plasmid # 12647; http://n2t.net/addgene:12647; RRID:Addgene_12647) (Brzovic *et al*., 2006).

### Protein expression

USP28 was expressed in *E. coli* strain MDS42. Ube1 (E1) and Ube2D1 (E2) were expressed in *E. coli* strain Rosetta2 (DE3). All other constructs were expressed in *E. coli* strain BL21(DE3). For proteins produced in *E. coli*, a starter culture was prepared in 5 mL of LB medium (Formedium, Hunstanton, Norfolk, England) with the appropriate antibiotic. After 14 h of incubation at 37°C, the starter culture was used to inoculate 500 mL of Terrific Broth (TB) autoinduction media with trace elements (Formedium, Hunstanton, Norfolk, England) supplemented with glycerol (8 g/L) and the appropriate antibiotic. Cultures were incubated briefly at 37°C until they reached and OD_600_ of 1.0, point after which the temperature was decreased to 18°C for overnight incubation. Cells were harvested by centrifugation at 4200xg for 45 minutes at 4°C and re-suspended in lysis buffer [50 mM HEPES (pH 7.5), 500 mM NaCl, 0.5 mM TCEP, 10% (v/v) glycerol, 10 mM imidazole]. Re-suspended pellets were flash frozen in liquid N_2_ and stored at -20°C until purification.

### Protein purification

Except ubiquitin, all constructs were initially purified by immobilised-metal affinity chromatography (IMAC). Cell pellets were lysed by sonication, and centrifuged (30000 x g, 45 min) to recover soluble material, which was filtered with a 0.45 μm sterile syringe filter (Sartorius). Filtered material was loaded to a 5 mL HiTrap IMAC HP column (GE Healhcare Biosciences) loaded with Ni^2+^ and pre-equilibrated with lysis buffer. The column was washed with 5 column volumes (CV) of lysis buffer followed by 5 CVs of IMAC wash buffer. Proteins were eluted from the column with IMAC elution buffer. After IMAC, Ube2D1 was immediately treated with SUMO-Specific protease 2 (SENP2) to cleave the His-sumo tag. Protein digestion was followed by a Reverse IMAC (R.IMAC) to remove the Sumo tag.

DTX3L and DTX3L^CS^ were subjected to MBP-affinity chromatography (following IMAC) by loading IMAC elutions to a 5 mL MBPTrap HP column (GE Healhcare Biosciences) equilibrated with SEC buffer. Column was washed with 4 CVs of SEC buffer and the proteins were recovered in MBP elution buffer. Subsequently, proteins were treated with TEV protease overnight (1:30 molar ratio, 4°C) to cleave purification tags, which were removed by R.IMAC.

After IMAC, Ube1, CFP-Ub and YFP-Ub were dialysed Anion Exchange Chromatography (AnEX) low salt buffer for at least 16 h at 4°C. After dialysis, proteins were loaded to a HiTrap Q HP XL column (GE Healhcare Biosciences). The column was washed with 4 CVs of AnEX low salt buffer and then connected to a BioRad system to run a gradient from low salt to high salt buffer at 2 mL/min. IMAC for USP28 was done by washing the column with IMAC buffer (4 CVs), followed by 4 CVs of ATP wash buffer. Immediately after, protein was washed with 4 CVs of IMAC Gradient 1 buffer, eluting the protein in a gradient from 100% IG1 buffer to 100% IMAC Gradient 2 buffer.

Size Exclusion Chromatography (SEC) was used as a final purification step for all proteins using a Superdex S75 16/600 or Superdex S200 16/600 column (depending on the MW of the target protein). Finally, positive SEC fractions were flash frozen with liquid N_2_ and stored at -70°C, hence storage SEC buffer also acts as a storage buffer. Buffers used for each the purification step are listed in **Supplementary Table S5**.

### Ub purification

Cell pellet of Ub was sonicated and the soluble fraction was recovered without filtration. Afterwards, soluble protein fraction was boiled at 80°C for 5 min. Protein aggregates were removed by centrifugation at 30000 x g (30 min, 4°C). Trichloro acetic acid (TCA) was added to the supernatant to precipitate proteins [final concentration of 5% (v/v)] and the mixture was incubated at room temperature for 10 min. After centrifugation (2000 rpm, 2 min), protein pellet was re-suspended in 15 mL of phosphate buffer saline (PBS). pH was adjusted to 7.4 to solubilise proteins, which were recovered by centrifugation at 2000 rpm for 2 min, 4°C. Supernatant was dialysed overnight at 4°C against Cation exchange (CatEX) low salt buffer. Following dialysis, protein was loaded to a 5 mL HiTrap SP column (GE Healhcare Biosciences) and then connected to a BioRad system. Proteins were eluted with a gradient from 100% low salt buffer to 100% CatEX high salt buffer. Ub-containing fractions were loaded to a Superdex S75 16/60 column for SEC. Protein was flash frozen with liquid N_2_ and stored at -70°C. Buffers used in each purification step are listed in **Supplementary Table S5**.

### Circular dichroism spectroscopy (CD)

DTX3L and DTX3L^CS^, were diluted to 0.1 mg/mL in phosphate buffer to decrease the absorbance generated by HEPES buffer. Storage buffer is diluted at the same ratio as the protein and used as a blank. The spectra were measured in triplicates from 190 to 280 nm. For the determination of protein melting temperature, spectra from 190 to 280 nm were measured throughout a temperature range of 22 to 80°C at an increase rate of 1°C/min. Measurements were recorded using a Chirascan™ CD spectrometer (Applied Photophysics) and data analysis was done using Chirascan software.

### Gel-based auto-ubiquitination assays

Ubiquitination reactions for DTX3L constructs were prepared in 50 μL volume by mixing E1 (Ube1; 0.4 μM), E2 (Ube2D1; 2 μM), E3 (DTX3L; 1 μM) and Ub (30 μM). Tris buffer (1 M, pH 7.5) containing ATP (40 mM), MgCl_2_ (100 mM) and DTT (40 mM) was added in a 1:20 (buffer to reaction volume) ratio to initiate ubiquitination. Upon ATP addition, the reaction was incubated at room temperature for up to 2 h. Auto-ubiquitination was quenched by adding 2x Laemmli buffer, and subsequently analysed with SDS-PAGE and PageBlue staining (Thermo Scientific). De-ubiquitination reactions were performed in a similar manner, with the addition of USP28 (45 μM) and the incubation time was extended to 4 h to ensure depletion of ATP.

### Ubiquitination FRET measurements

A 30x ubiquitination mix was prepared and dispensed with Echo acoustic dispenser (Labcyte). Final enzyme concentrations in the reaction were: E1 (Ube1; 12 nM), E2 (Ube2D1; 180 nM), E3 (DTX3L; 500 nM), CFP-Ub (25 nM) and YFP-Ub (350 nM). To initiate the reaction, ubiquitination buffer [5 mM Tris (pH 7.5), 200 μM ATP, 0.5 mM MgCl_2_, 200 μM DTT, (final concentrations)] was added to the plate using Mantis liquid dispenser (Formulatrix). Measurements were performed using the multimode microplate reader TECAN Infinite M1000 PRO or TECAN Spark using a fluorescence top reading modality. Excitation wavelength was set to 410 nm (20 nm bandwidth). The emission reading wavelength was set to 477 nm and 527 nm, both at 10 nm bandwidth. Reading gain was set manually to 80% throughout the study. The number of flashes was set to 50 at 400 Hz frequency. Integration time was 20 μs and 44 ms were set for the settling time. Z-position of the reader was fixed to 24019 μm. For the measurements we used 384-well polypropylene, black, flat-bottom plates (Greiner). Except for the buffer optimisation tests, all measurements were performed in reaction buffer [10 mM HEPES (pH7.5)] in 20 μL. Except for the optimisation tests, reactions were measured right after the addition of ubiquitination buffer and after an incubation of 90 min at room temperature.

### Signal verification and statistical validation for ubiquitination assay

Experiments were performed in 384-well polypropylene, black, flat-bottom plates (Greiner). rFRET of reactions with DTX3L was compared to the rFRET of reactions with DTX3L^CS^ to confirm that the signal was caused by the formation of poly-Ub chains. For each condition, 4 data points were collected in a 20 μL reaction volume.

For statistical validation of the assay we calculated the Z’ factor (Zhang, Chung and Oldenburg, 1999). Additionally, we calculated the signal to background ratio by dividing the average signal by the average background. The signal to noise ratio was calculated by dividing the specific signal by the standard deviation, we defined specific signal as the difference between the maximum signal and the minimum signal. FRET ubiquitination reactions were mixed in 20 μL and the fluorescence was measured in presence and absence of 50 mM EDTA, with 880 replicates for each condition in an interleaved plate-format.

### Compound screening

As a way to validate the functionality of the assay, we screened 2428 compounds (at 100 nM final concentration) from three libraries of the NCI against the ubiquitination reaction with DTX3L. Compounds were transferred to 384-well plates containing E1 (UBE1; 12 nM), E2 (Ube2D1; 180 nM), E3 (DTX3L, 500 nM), Ub_Don_ (25 nM) and Ub_Acc_ (350 nM). To initiate the reaction, ubiquitination buffer [5 mM Tris (pH 7.5), 200 μM ATP, 0.5 mM MgCl_2_, 200 μM DTT, (final concentrations)] was added to the plate using Mantis liquid dispenser (Formulatrix). Reactions were filled to 20 μL with 10 mM HEPES (pH 7.5) and buffer supplemented with 50 mM EDTA was used as a positive control. During the screening, the rFRET was calculated at two excitation wavelengths (410 nm and 430 nm). MarvinSketch (Chemaxon) was used for drawing chemical structures.

### Differential scanning fluorimetry (DSF)

Measurements were performed using Prometheus Nt.48 NanoDSF (NanoTemper) coupled with standard capillaries. DTX3L was diluted in 10 mM HEPES (pH 7.5) to 4 mg/mL and determination of the melting temperature was done in the presence and absence of 100 μM of selected compounds from the screening. Melting curves were recorded from 20-95°C, with a temperature increase of 1°C/min. Data analysis and calculation of the melting temperature (Tm) was done in GraphPad Prism 9 by fitting the data to the Boltzmann sigmoidal equation. Every condition was tested in triplicates.

### De-ubiquitination FRET measurements

To study the hydrolysis of the poly-Ub chains by USP28, ubiquitination reactions were performed as described above but incubated for 150 min to ensure maximal FRET signal. Subsequently, EDTA (50 mM final concentration) was added to stop reaction from progressing and immediately after, USP28 (50 nM) was added. The final reaction volume was 20 μL. The reaction was incubated for up to 1 h with reading intervals of 15 min. A decrease in the rFRET was interpreted as chain hydrolysis by USP28.

Verification of the signal was done by comparing the rFRET of reaction with USP28 to reactions with USP28^C174A^. Statistical validation of the hydrolysis was done in a similar way as for DTX3L, with the variation that absence of USP28 (maximum rFRET) was considered as positive control and presence of USP28 (minimum rFRET) was considered as negative control. The

## Acknowledgments

Biocenter Oulu Structural Biology core facility, member of Biocenter Finland, Instruct-ERIC Centre Finland and FINStruct, as well as of “Proteomics and Protein Analysis” and Sequencing core facilities are gratefully acknowledged.

## Author contributions

C.V.-R., Y.A., L.L. conceptualization; C.V.-R., I.S. investigation; C.V.-R., L.L. writing-original draft; C.V.-R., I.S., Y.A., L.L. writing-reviewing and editing; L.L. funding acquisition; L.L. supervision.

## Additional Information

The authors declare no competing interests.

