## Supplementary figures and tables for "Discovery of DTX3L inhibitors through a homogeneous FRET-based assay that monitors formation and removal of poly-ubiquitin chains"

[&]Current address: Department of Genome Sciences, University of Washington, Seattle, USA

### Contents

Figure S1. Amino acid sequence alignment between the original fluorescent constructs and the final constructs used in the assay.
Figure S2. Static light scattering and molar mass distribution for the CFP-Ub and YFP-Ub.
Figure S3. CD spectra of the mutant DTX3L compared to the WT.
Figure S4. Gel-based ubiquitination assay of DTX3L and DTX3L$^{CS}$.
Figure S5. Time-dependent rFRET of the initial trials of the ubiquitination reaction with DTX3LCS.
Figure S6. Individual charts of the optimisation stage.
Figure S7. Time-dependent plot for the last optimisation step of the assay conditions.
Table S1. Statistical parameters of the plate for the ubiquitination assay with DTX3L.
Figure S8. Gel-based ubiquitination assay of DTX3L in presence of the compounds identified in the screening stage.
Table S2. Parameters obtained from nanoDSF for DTX3L with 100 µM compound.
Table S3. Statistical parameters of the plate for the de-ubiquitination assay.
Table S4. Expression constructs used in recombinant protein production.
Table S5. Composition of buffers used in protein purification.

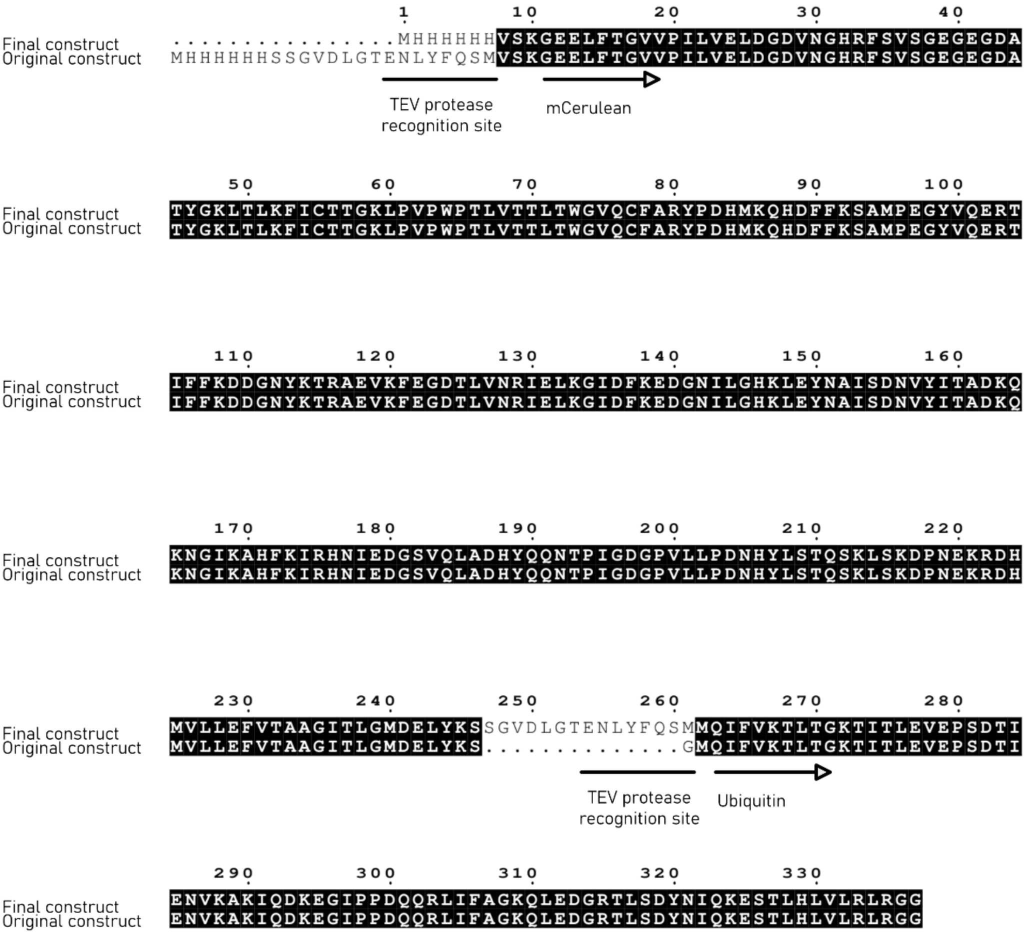


**Figure S1.** Amino acid sequence alignment of the constructs for Ub-CFP fusion proteins. The upper row corresponds to the construct used in the assay consisting of a 6X-His tag, Ub, linker including a TEV cleavage site and CFP. The lower row corresponds to the construct that generated inclusion bodies due to the lack of the linker.

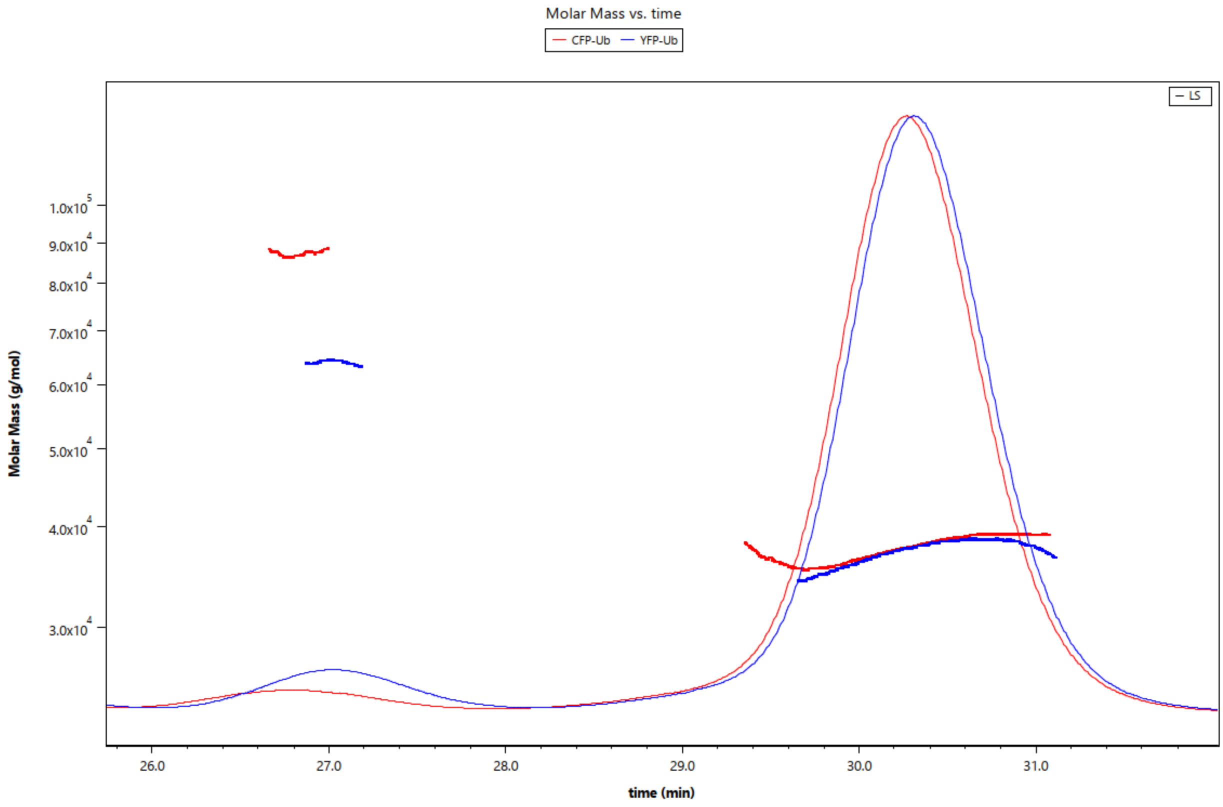


**Figure S2.** Static light scattering and molar mass distribution for the CFP-Ub and YFP-Ub. The purified fusion construct runs as a monomer as calculated from the refractive index of the sample. The calculated molecular weight for YFP-Ub and CFP-Ub is 38 kDa and the experimentally determined value is 37 kDa.

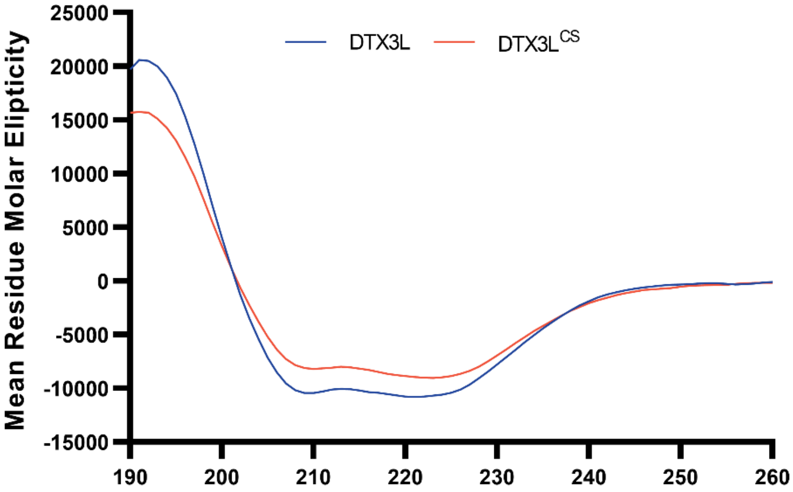


**Figure S3.** CD spectra of the WT DTX3L compared to DTX3L$^{CS}$. Spectra indicates that mutation in the RING finger of DTX3L does not alter to a great extent the folded state of the protein.

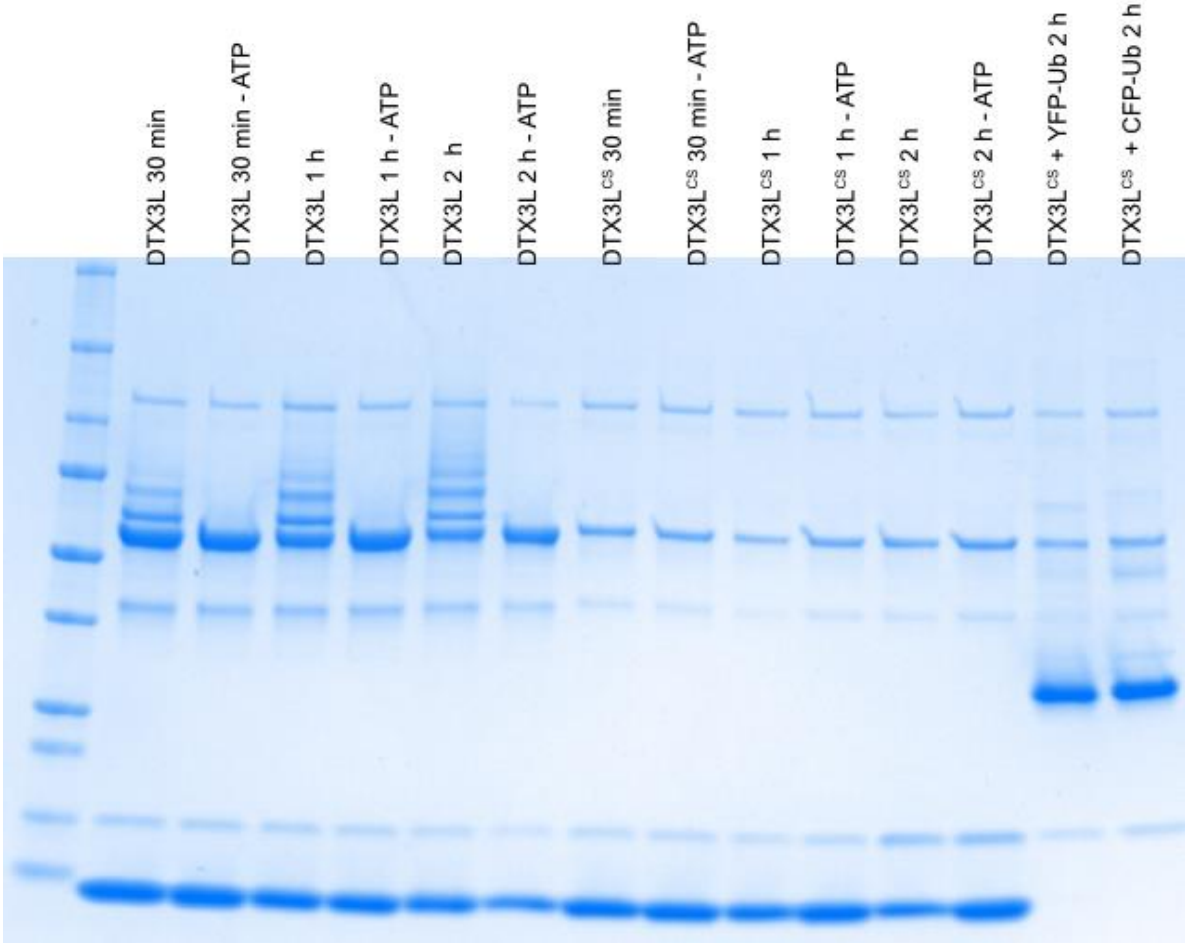


**Figure S4.** Gel-based ubiquitination assay of DTX3L (lanes 1-7) and DTX3L$^{CS}$ (lanes 8-13) show that DTX3L$^{CS}$ is unable to undergo auto-ubiquitination.

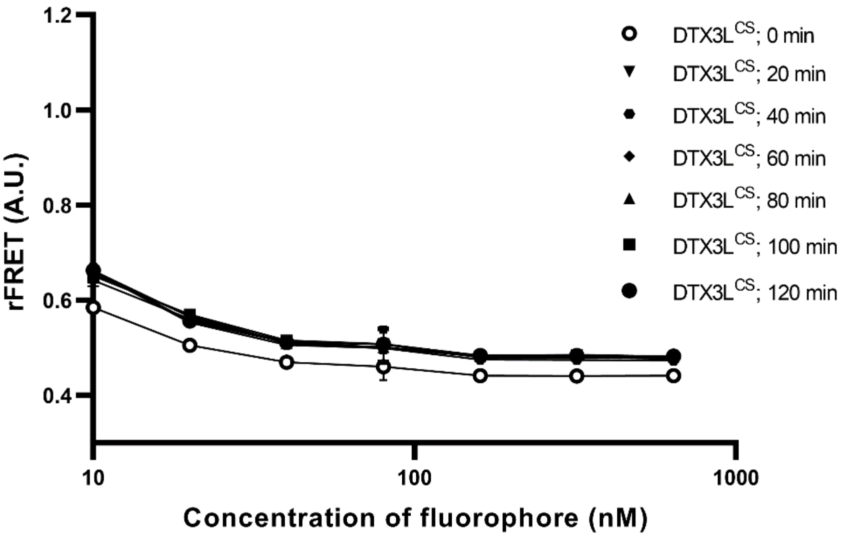


**Figure S5.** Time-dependent rFRET of the initial trials of the ubiquitination reaction with DTX3L$^{CS}$ confirm the lack of ligase activity also in the context of the FRET-based assay as compared to data shown in **Figure 2A**. Data shown are mean + SD, n=4.

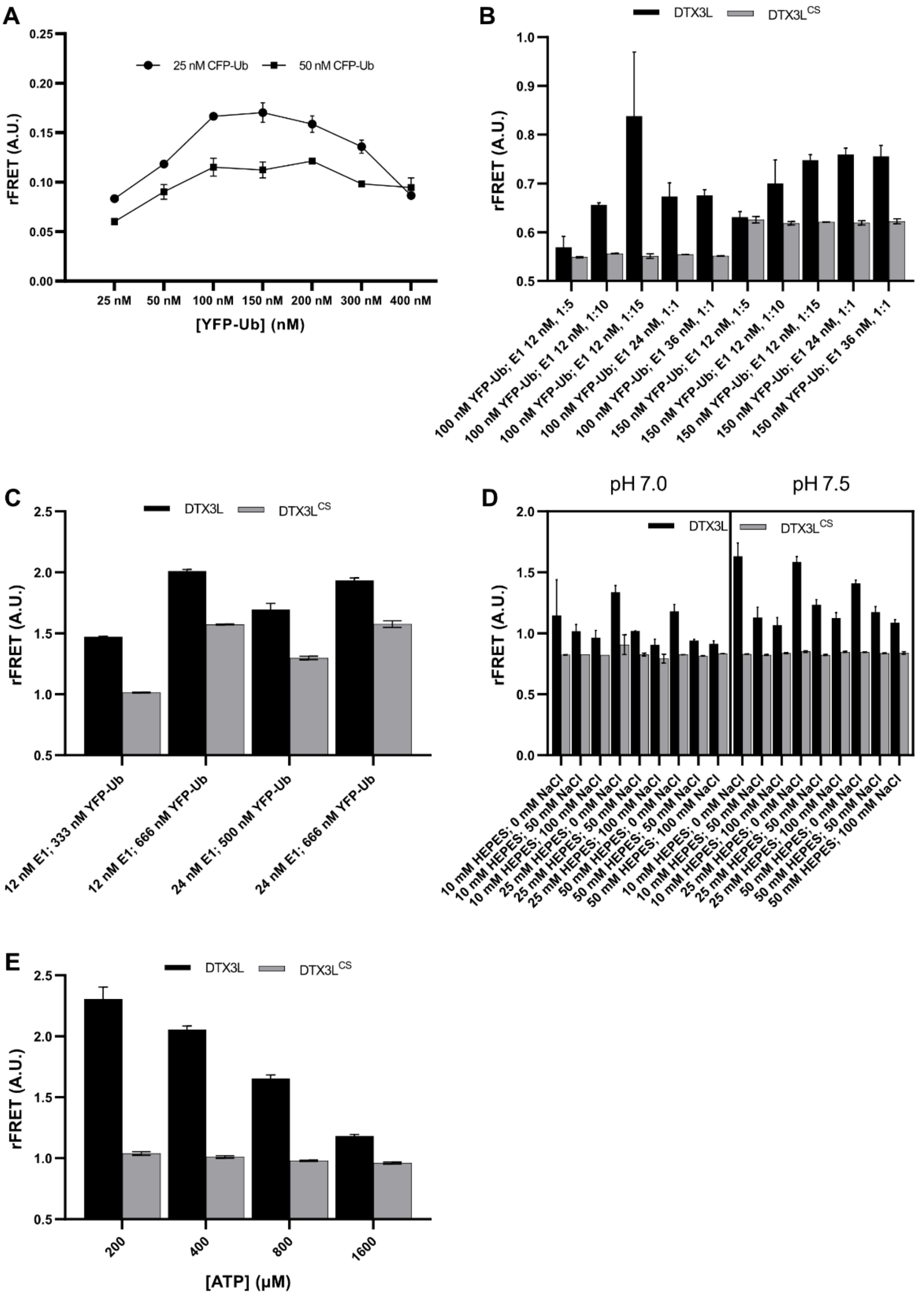
A
rFRET (A.U.)
25 nM CFP-Ub
50 nM CFP-Ub
[YFP-Ub] (nM)
25 nM
50 nM
100 nM
150 nM
200 nM
300 nM
400 nM
B
DTX3L
DTX3L$^{CS}$
100 nM YFP-Ub; E1 12 nM, 1:5
100 nM YFP-Ub; E1 12 nM, 1:10
100 nM YFP-Ub; E1 12 nM, 1:15
100 nM YFP-Ub; E1 24 nM, 1:1
100 nM YFP-Ub; E1 36 nM, 1:1
150 nM YFP-Ub; E1 12 nM, 1:5
150 nM YFP-Ub; E1 12 nM, 1:10
150 nM YFP-Ub; E1 12 nM, 1:15
150 nM YFP-Ub; E1 24 nM, 1:1
150 nM YFP-Ub; E1 36 nM, 1:1
C
12 nM E1; 333 nM YFP-Ub
12 nM E1; 666 nM YFP-Ub
24 nM E1; 500 nM YFP-Ub
24 nM E1; 666 nM YFP-Ub
D
pH 7.0
pH 7.5
10 mM HEPES; 0 mM NaCl
10 mM HEPES; 50 mM NaCl
10 mM HEPES; 100 mM NaCl
25 mM HEPES; 0 mM NaCl
25 mM HEPES; 50 mM NaCl
25 mM HEPES; 100 mM NaCl
50 mM HEPES; 0 mM NaCl
50 mM HEPES; 50 mM NaCl
50 mM HEPES; 100 mM NaCl
E
[ATP] (µM)
200
400
800
1600

**Figure S6.** Individual charts of the optimisation stages. A) Titration of CFP-Ub at two concentrations (25 and 50 nM) with different concentrations of YFP-Ub. Reactions also contained Ube1 (12 nM), Ube2D1 (60 nM) and DTX3L (500 nM). B) rFRET of ubiquitination reactions at two concentrations of YFP-Ub (100 and 150 nM) varying the concentrations of Ube1 (12, 24 and 36 nM) and the ratio of Ube1:Ube2D1. All reactions contained CFP-Ub (25 nM) and DTX3L (500 nM) C) rFRET of ubiquitination reactions at varying concentrations of YFP-Ub (333, 500 and 666 nM) and Ube1 (12 and 24 nM). All reactions contained CFP-Ub (25 nM), DTX3L (500 nM) and Ube2D1 (1:15 Ube1:Ube2D1 ratio). C) rFRET of ubiquitination reactions at pH 7.0 and 7.5 and varying concentrations of HEPES and NaCl. E) rFERT of ubiquitination reactions at varying concentrations of ATP. Reactions contained CFP-Ub (25 nM), YFP-Ub (350 nM), Ube1 (12 nM), Ube2D1 (180 nM) and DTX3L (500 nM). Reactions of panels A-D contained 2 mM ATP. Data shown in panels C and E correspond to mean + SD, n=3. Data shown in panels A-B, D correspond to mean ± SD, n=2.

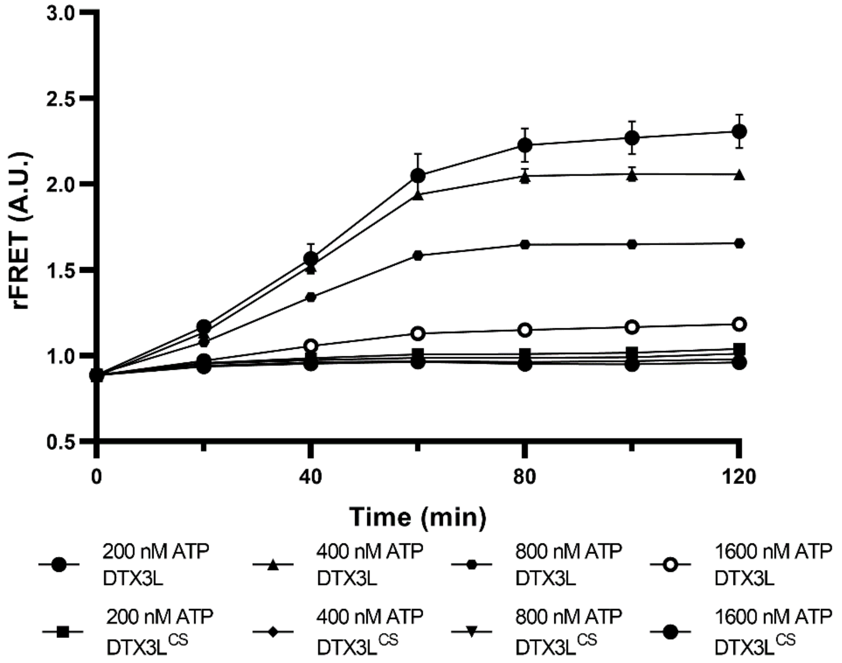


**Figure S7.** Time-dependent plot for the last optimisation step of the assay conditions in which different conce. Data shown are mean + SD, n=4.

**Table S1. Statistical parameters of the plate for the ubiquitination assay of DTX3L.**

| | **Day 1** | **Day 2** | **Day 3** | | | **Mean** | **SD** | **CV (%)** |
|---|---|---|---|---|---|---|---|---|
| | | | **Plate 1** | **Plate 2** | **Plate 3** | | | |
| **Maximum** | 1.96 | 2.6 | 2.55 | 2.65 | 2.32 | 2.42 | 0.28 | 11.7 |
| **Minimum** | 0.76 | 0.95 | 0.84 | 0.89 | 0.82 | 0.85 | 0.07 | 8.3 |
| **Z'** | 0.7 | 0.73 | 0.78 | 0.85 | 0.76 | 0.76 | 0.06 | 7.5 |
| **S/N** | 155 | 67 | 99 | 139 | 130 | 118 | 35 | 29.9 |
| **S/B** | 2.6 | 2.7 | 3.0 | 3 | 2.8 | 2.8 | 0.18 | 6.4 |
| **Day-to-day** | | | | | | | | 1.02-4.86 |
| **Plate-to-plate** | | | | | | | | 1.42-4.67 |

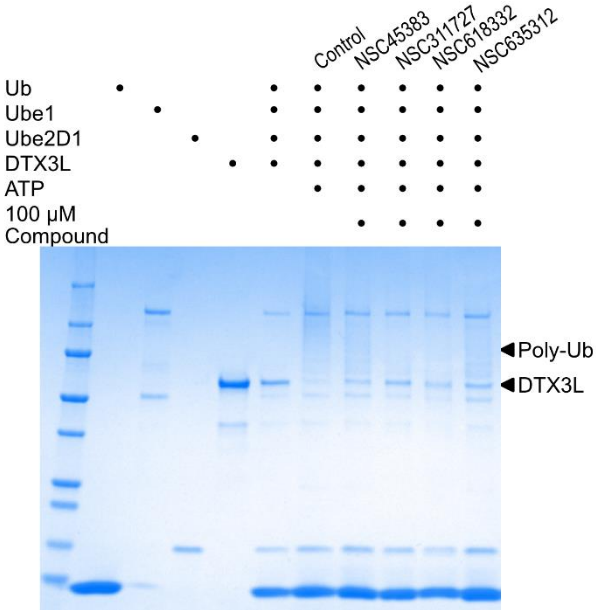


**Figure S8.** Gel-based ubiquitination assay of the compounds identified in the screening stage seem to inhibit auto-ubiquitination of DTX3L based on the characteristic smear of poly-ub chains.

**Table S2. Parameters obtained from nanoDSF for DTX3L with 100 µM compounds.**

| **Treatment** | **Tm (°C)** | **Onset for scattering (°C)** | **Difference in Tm (°C)** |
|---|---|---|---|
| **DMSO** | 41.7 ± 0.04 | 40.3 ± 0.39 | |
| **NSC45383** | 47.0 ± 0.13 | 40.4 ± 0.65 | 5.2 |
| **NSC311727** | 49.6 ± 0.10 | 36.7 ± 0.78 | 7.9 |
| **NSC618332** | 48.7 ± 0.03 | 32.9 ± 0.83 | 7 |
| **NSC635312** | 48.8 ± 0.02 | 39.1 ± 0.41 | 7.1 |

**Table S3. Statistical parameters of the plate for the de-ubiquitination assay.**

| | Day 1 | Day 2 | | Mean | SD | CV (%) |
|---|---|---|---|---|---|---|
| | | Plate 1 | Plate 2 | | | |
| **Maximum** | 2.2 | 2.1 | 2..1 | 2.1 | 0.04 | 2.2 |
| **Minimum** | 1.6 | 1.7 | 1.7 | 1.6 | 0.04 | 2.8 |
| **Z'** | 0.69 | 0.72 | 0.73 | 0.71 | 0.01 | 2.4 |
| **S/N** | 21.1 | 24.8 | 24.9 | 23.6 | 1.8 | 7.5 |
| **S/B** | 1.4 | 1.2 | 1.2 | 1.3 | 0.1 | 7.4 |
| **Day-to-day** | | | | | | 0.85 – 1.7 |
| **Plate-to-plate** | | | | | | 0.85 – 0.92 |

**Table S4. Expression constructs used in recombinant protein purification.**

| Construct | UniProt ID | Amino acid boundaries | Vector | Expression system |
|---|---|---|---|---|
| Uba1 (Addgene #34965) | P22314 | 1-1058 | pET21d | *E. coli* |
| Ube2D1 (Addgene #61081) | P51668 | 1-147 | pETSUMO | *E. coli* |
| Ubc (Addgene #12647) | P0CG48 | 1-76 | pET15 | *E. coli* |
| CFP-Ubc | P0CG48 | 1-76 | pNIC-CFP | *E. coli* |
| YFP-Ubc | P0CG48 | 1-76 | pNIC-YFP | *E. coli* |
| DTX3L FL | Q8TDB6 | 1-740 | pFastBac1 His-MBP (Addgene #30116) | Sf21 insect cells |
| DTX3L D3RD | Q8TDB6 | 230-740 | pNIC-MBP | *E. coli* |
| DTX3L D3RD CS | Q8TDB6 | 230-740 (C561S, C564S) | pNIC-MBP | *E. coli* |
| USP28 | Q96RU2 | 1-1077 | pMJS162 | *E. coli* |

**Table S5. Composition of buffers used in protein purification.**

| Buffer | Composition |
|---|---|
| Lysis | 50 mM HEPES (pH 7.5), 500 mM NaCl, 0.5 mM TCEP, 10% (v/v) glycerol, 10 mM imidazole |
| IMAC wash | 50 mM HEPES (pH 7.5), 500 mM NaCl, 0.5 mM TCEP, 10% (v/v) glycerol, 25 mM imidazole |
| IMAC elution | 50 mM HEPES (pH 7.5), 500 mM NaCl, 0.5 mM TCEP, 10% (v/v) glycerol, 300 mM imidazole |
| SEC | 30 mM HEPES (pH 7.5), 350 mM NaCl, 0.5 mM TCEP, 10% (v/v) glycerol |
| Reverse IMAC wash | 50 mM HEPES (pH 7.5), 500 mM NaCl, 0.5 mM TCEP, 10% (v/v) glycerol, 50 mM imidazole |
| MBP elution | 30 mM HEPES (pH 7.5), 350 mM NaCl, 0.5 mM TCEP, 10% (v/v) glycerol, 10 mM maltose |
| UBA1 AnEX low salt | 20 mM Tris (pH 8), 50 mM NaCl, 0.5 mM TCEP, 10% (v/v) glycerol |
| UBA1 AnEX high salt | 20 mM Tris (pH 8), 500 mM NaCl, 0.5 mM TCEP, 10% (v/v) glycerol |
| Fluorescent Ubc AnEX low salt | 50 mM HEPES (pH 7.5), 50 mM NaCl, 0.5 mM TCEP |
| Fluorescent Ubc AnEX high salt | 50 mM HEPES (pH 7.5), 500 mM NaCl, 0.5 mM TCEP |
| USP28 ATP wash | 50 mM HEPES (pH7.5), 2 mM ATP, 5 mM $MgCl_2$, 2 mM DTT |
| USP28 IMAC gradient 1 (IG1) | 50 mM HEPES (pH 7.5), 500 mM NaCl, 0.5 mM TCEP, 10% (v/v) glycerol |
| USP28 IMAC gradient 2 (IG2) | 50 mM HEPES (pH 7.5), 500 mM NaCl, 0.5 mM TCEP, 10% (v/v) glycerol, 10 mM imidazole |
| Ub lysis | 50 mM HEPES (pH 7.5) |
| Ub CatEX low salt | 50 mM Ammonium acetate (pH 4.5) |
| Ub CatEX high salt | 50 mM Ammonium acetate (pH 4.5), 500 mM NaCl |
| Ub SEC buffer | 50 mM HEPES (pH 7.5) |
